# Disruption of lamina-associated genome organization activates neuronal gene programs in H3.3K27M DIPG

**DOI:** 10.64898/2026.09.21.753114

**Authors:** Sajad Hamid Ahanger, Mitchel A. Cole, Eugene Gil, Evan R. Semenza, Joanna J. Phillips, Daniel A. Lim

## Abstract

Diffuse intrinsic pontine glioma (DIPG) is a fatal pediatric brain tumor frequently driven by a histone H3 mutation that substitutes lysine 27 with methionine (H3K27M). While H3K27M is known to cause a global depletion of the repressive histone mark H3K27me3, how this epigenetic disruption contributes to oncogenic transcriptional reprogramming remains incompletely understood. Here, we show that H3.3K27M DIPG cells exhibit widespread disruption of lamina-associated domain (LAD) organization across primary tumors and patient-derived cell lines. H3.3K27M was enriched at genomic regions that lost lamina association, which acquired increased H3K27ac and showed activation of neuronal and oncogenic gene programs. Notably, LAD loss was more strongly associated with transcriptional activation than H3K27me3 depletion, linking altered LAD organization to the transcriptional consequences of H3.3K27M. Finally, expression of H3.3K27M in human embryonic stem cell-derived neural precursor cells (NPCs) was sufficient to induce LAD remodeling, predominantly through loss of lamina association, leading to activation of neuronal gene programs. Together, these findings identify disruption of lamina-associated genome architecture as a key mechanism by which the H3.3K27M oncohistone drives oncogenic transcriptional reprogramming in DIPG.

## Introduction

Diffuse intrinsic pontine glioma (DIPG) is the leading cause of brain tumor-related death in children with no effective treatment options(1). Over 80% of DIPGs harbor a lysine-to-methionine substitution at position 27 of histone H3 (H3K27M), most commonly in the *H3F3A* gene encoding the H3.3 variant(2–4). This mutation exerts a dominant inhibitory effect on the enzymatic activity of enhancer of zeste homolog 2 (EZH2), the catalytic subunit of the polycomb repressive complex 2 (PRC2), leading to a global reduction in histone H3 lysine 27 trimethylation (H3K27me3) – a key epigenetic mark associated with transcriptional repression(5,6). While the loss of H3K27me3 is widely considered central to DIPG pathogenesis, its direct relationship to the oncogenic transcriptional program remains weak(7,8). Furthermore, H3.3K27M-mutant cells exhibit epigenetic alterations beyond PRC2 inhibition(9–13), suggesting that the oncohistone drives tumorigenesis through additional unknown mechanisms.

Altered nuclear morphology is a hallmark of many cancers and is often associated with changes in heterochromatin organization(14). In particular, H3.3K27M expression has been linked to reduced nuclear size and circularity(15), suggesting that this oncohistone may perturb spatial genome architecture. Yet, how the H3.3K27M oncohistone influences subnuclear compartments remains unknown.

In mammalian cells, nearly half of the genome is anchored to the nuclear periphery through lamina-associated domains (LADs)(16). LADs are large (10 Kb – 10 Mb), heterochromatic regions typically enriched for repressive histone modifications, including H3K27me3(17). LADs help partition the genome into spatial compartments within the nucleus and act as dynamic regulators of gene expression, thereby maintaining cell-type–specific transcriptional programs(18). Despite their fundamental role in gene regulation, it remains unknown whether LAD architecture is disrupted in cancer, particularly in the context of H3K27M-driven epigenomic dysregulation.

In this study, we investigated how the H3.3K27M oncohistone alters higher-order genome organization in DIPG. Using primary tumor specimens, patient-derived cell lines, and human embryonic stem cell-based models, we show that H3.3K27M expression disrupts LAD architecture. This spatial reorganization of the genome is closely associated with activation of gene expression programs implicated in DIPG pathogenesis. Together, our findings uncover a previously unrecognized mechanism by which the H3.3K27M oncohistone drives transcriptional reprogramming through disruption of spatial 3D genome architecture.

## Results

### LAD architecture is disrupted in primary DIPG tumors harboring H3.3K27M

To examine the LAD organization in primary DIPG tumors, we performed fluorescence-activated nuclei sorting (FANS) using an H3K27M-specific antibody to isolate H3.3K27M-positive (H3K27M-Pos) and H3.3K27M-negative (H3K27M-Neg) nuclei from three post-mortem DIPG specimens (**Fig. 1a**). RNA-seq of sorted nuclei revealed that samples clustered by H3 mutation status, with G2/M and oligodendrocyte precursor cell (OPC) states(19) predominating within each tumor sample (**Fig. 1b, Extended Data Fig. 1a**). We then profiled H3K27me3 by CUT&RUN and LADs by LaminB1 GO-CaRT(20,21). To prevent off-target cleavage from pAG-MNase binding to the H3K27M antibody used during FANS, nuclei were pre-incubated with monovalent Fab fragments prior to CUT&RUN and GO-CaRT as previously described (**Fig. 1a**)(22). Consistent with the known effects of the mutation, H3K27M-Pos nuclei exhibited global loss of H3K27me3 relative to H3K27M-Neg nuclei (**Fig. 1c**), validating the specificity of our FANS-based approach.

**Figure 1.**
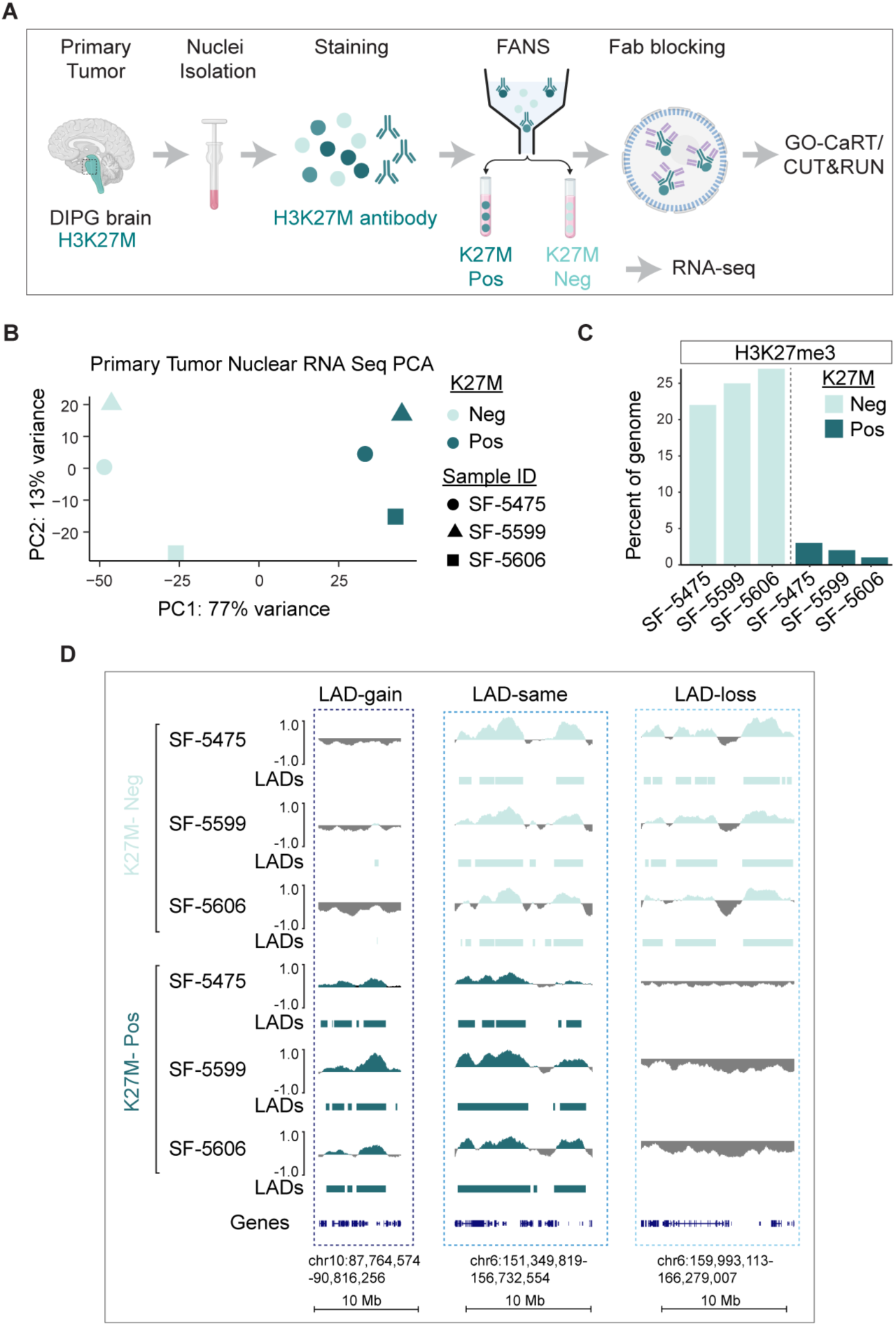
H3K27M-positive nuclei from primary DIPG tumor samples have altered LAD profiles from that of paired H3K27M negative nuclei. **A,** Experimental overview of FANS to separate H3K27M-positive nuclei and H3K27M-negative nuclei in primary DIPG tumor samples. Sorted nuclei were processed for RNA-seq or GO-CaRT/CUT&RUN with prior Fab blocking **B,** Principal Component Analysis of RNA-seq from H3K27M-positive nuclei and H3K27M-negative nuclei from primary tumors. Colors indicate H3K27M status and symbols indicate tumor identity **C,** Percentage of analyzed genome (chr1-chr22, chrX with blacklisted regions removed) under H3K27me3 peaks in each population. **D,** Representative genome browser views of LADs gained in H3K27M-positive nuclei, same in both H3K27M-positive and H3K27M-negative nuclei, and LADs lost in H3K27M-positive nuclei in hg38 coordinates. For each tumor-genotype combination, the top track represents log2(LMNB1/IgG) signal and bottom track represent LAD calls for each locus.

Genome-wide LAD coverage was broadly similar between H3K27M-Pos and H3K27M-Neg nuclei within each tumor, ranging from 35–48% (**Extended Data Fig. 1b**). In both populations, LADs retained characteristic heterochromatic features, including lower gene and SINE density, enrichment of LINE elements, and higher levels of H3K9me2 – a histone modification typically enriched in LADs (**Extended Data Fig. 1c-d**).

Despite similar overall LAD coverage, PCA of genome-wide LaminB1 signal separated H3K27M-Pos from H3K27M-Neg nuclei, indicating redistribution of lamina-associated chromatin in H3K27M-mutant cells (**Extended Data Fig. 1e**). While 29-37% (859-1095 Mb) of the genome remained associated with the lamina in both populations (LAD-same), 15-17% (434-491 Mb) underwent LAD reorganization in H3K27M-Pos nuclei, comprising regions of LAD loss (5-7%, 136-210 Mb) and LAD gain (8-11%, 224-326 Mb) relative to H3K27M-Neg nuclei (**Fig. 1d; Extended Data Fig. 2a**). Importantly, these LAD alterations were highly recurrent across tumors, with 242 Mb (8% genome) of LAD-gain and 137 Mb (5% genome) of LAD-loss regions shared across at least two of three primary tumors (**Extended Data Fig. 2b**). Regions undergoing LAD reorganization also showed corresponding alterations in H3K9me2 (**Extended Data Fig. 2c-d**). Thus, H3.3K27M mutation is associated with widespread disruption of LAD architecture in primary DIPG tumors.

### LAD loss, rather than H3K27me3 depletion, is associated with gene activation in H3.3K27M DIPG

We next performed gene ontology (GO) analysis of genes located within differential LAD regions. Genes residing in LAD-loss regions (n = 254) were enriched for developmental and oncogenic pathways previously implicated in DIPG biology(4,23,24), including processes related to synapse assembly, axonogenesis, and cell adhesion (**Extended Data Fig. 3a**). In contrast, genes within LAD-gain regions (n = 1596) were enriched for immune-related processes, such as natural killer cell activation and positive regulation of STAT protein phosphorylation (**Extended Data Fig. 3b)**.

To determine the transcriptional consequences of LAD reorganization in DIPG, we compared gene expression profiles between H3K27M-Pos and H3K27M-Neg nuclei using RNA-seq. Consistent with the repressive nature of the nuclear lamina, genes located within LADs were expressed at significantly lower levels than genes in non-LAD regions across all three primary tumor samples, regardless of H3K27M status (**Extended Data Fig. 3c**).

We next asked whether transcriptional activation in H3K27M-Pos nuclei was more closely associated with LAD loss or H3K27me3 depletion. Genes within LAD-loss regions showed significantly increased expression in H3K27M-Pos nuclei across all three tumors (**Fig. 2a**). In contrast, genes within H3K27me3-loss regions exhibited a modest increase in gene expression (**Fig. 2b**). Locus-specific analyses further illustrated the stronger association between lamina detachment and gene activation (**Fig. 2c, Extended Data Fig. 3d**). For instance, DIPG-associated genes *PTPRZ1* and *PTN*, as well as the neuronal genes *ADGRL3* and *LSAMP*, resided within LAD-loss regions and showed increased expression in H3K27M-Pos nuclei. In contrast, *PTGFR* and *ADGRL4*, which reside in regions that undergo pronounced H3K27me3 depletion but remain lamina associated (H3K27me3-loss/LAD-same), displayed no change in their low level of expression. Conversely, *PTPRC*, located within a LAD-gain region, showed reduced expression in H3K27M-Pos nuclei despite the absence of H3K27me3 enrichment in both H3K27M-Pos and H3K27M-Neg nuclei. These results indicate that transcriptional changes in H3.3K27M DIPG are more closely associated with altered lamina association than with H3K27me3 depletion alone, with LAD loss linked to the activation of neuronal and developmental gene programs.

**Figure 2.**
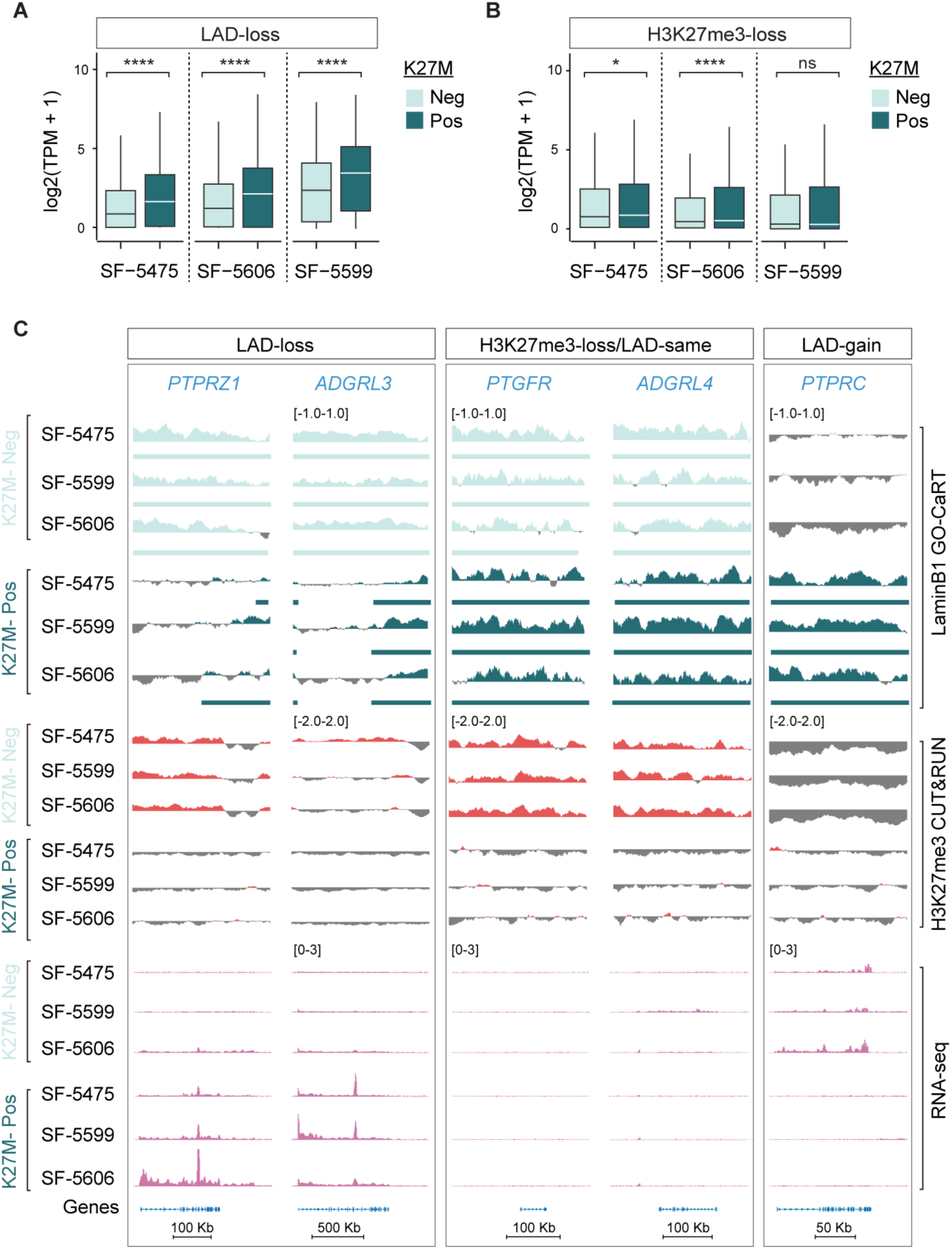
LAD loss regions exhibit greater transcriptional upregulation than H3K27me3 loss regions in primary DIPG tumors. **A, B** Box plots of gene expression in log2(TPM + 1) of genes in LAD-loss regions (**A**) or with H3K27me3 loss at promoters (**B)**, in H3K27M-positive and H3K27M-negative paired nuclei from primary DIPG tumors. Significance was assessed by unpaired Wilcoxon rank-sum test comparing H3K27M-positive with H3K27M-negative nuclei within each tumor; ns: p > 0.05; *: p ≤ 0.05; ****: p ≤ 1 × 10⁻⁴. **C,** Representative genome browser views of LAD-loss loci (*PTPRZ1*, *ADGRL3*), H3K27me3-loss loci with unchanged LAD status (*PTGFR*, *ADGRL4*), and LAD-gain locus (*PTPRC*). Within each assay, tracks are ordered H3K27M-negative (upper three) then H3K27M-positive (lower three) for the SF-5475, SF-5599, and SF-5606. LaminB1 GO-CaRT and H3K27me3 CUT&RUN signal tracks show log2(signal/IgG). Bars beneath the LaminB1 GO-CaRT tracks indicate LADs. RNA-seq coverage is shown in bins per million. Gene annotations and genomic scale are shown below.

### H3.3K27M DIPG cell lines exhibit widespread reorganization of facultative LADs

We next performed LaminB1 GO-CaRT to generate LAD profiles in three well-characterized patient-derived H3.3K27M DIPG cell lines: SU-DIPG-XIII, SU-DIPG-XVII, and SU-DIPG-XXIV (**Fig. 3a**). For comparison, LAD profiles were generated in two H3.3 wild-type (WT) glioma lines – VUMC-DIPG10 and pediatric cortical glioblastoma (pcGBM2). Each cell line was analyzed in duplicate, and next-generation sequencing (NGS) read coverage demonstrated high reproducibility between replicates (Spearman = 0.89 – 0.95, **Extended Data Fig. 4a**). Across cell types, LADs covered 34–48% of the genome and displayed canonical LAD features, including low gene and SINE density, and enrichment of LINE elements (**Extended Fig. 4b,c**), confirming that both H3.3WT and H3.3K27M DIPG cell lines form typical LADs.

**Figure 3.**
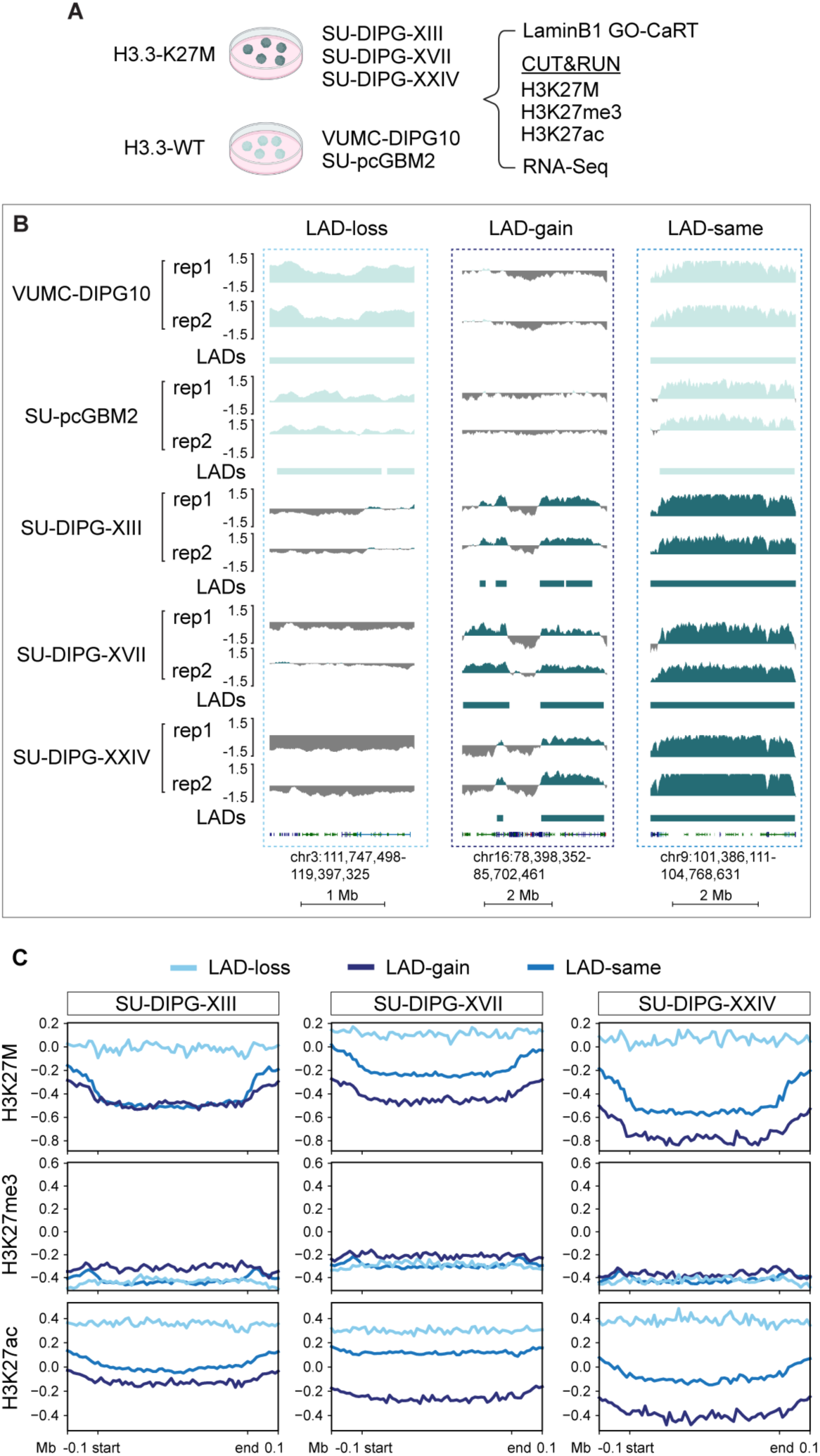
H3K27M is enriched in LAD loss regions in patient derived H3K27M-positive DIPG cell lines. **A,** Experimental overview of patient derived H3K27M-positive DIPG cell lines (SU-DIPG-XIII, SU-DIPG-XVII, SU-DIPG-XXIV) and H3WT pediatric glioma cell lines (VUMC-DIPG10 and SU-pcGBM2). **B,** Representative genome browser of LAD-loss, LAD-gain, and LAD-same regions of H3K27M-positive DIPG cell lines relative to H3WT glioma controls. Two replicates LMNB1 signal tracks presented as log2(LMNB1/IgG) with LAD calls displayed beneath signal tracks. **C,** Average H3K27M, H3K27me3, and H3K27ac signal profiles over LAD-loss, LAD-gain, and LAD-same loci. LAD-loss and LAD-gain regions were common to at least two H3K27M positive cell lines and absent in both WT controls. LAD-same were regions present in all 5 cell lines. The same regions were used for all three H3K27M-positive cell lines. Regions were scaled to 500 kb and flanked by 100 kb upstream and downstream; log2(signal/IgG) was averaged across all domains in each class in scaled 10-kb bins, and domains shorter than the 10-kb bin size were excluded. Columns show SU-DIPG-XIII, SU-DIPG-XVII, and SU-DIPG-XXIV. Rows show H3K27M, H3K27me3, and H3K27ac, signal.

To determine whether H3.3K27M DIPG cell lines exhibit LAD alterations similar to primary tumors, we compared LAD profiles between H3.3K27M and H3.3WT lines. PCA of genome-wide LaminB1 signal clearly separated H3.3K27M from H3.3WT samples, with the H3.3K27M cell lines occupying similar PCA space as primary tumors (**Extended Data Fig. 4d**). Approximately 22% of the genome (648 Mb) comprised LADs shared between H3.3K27M and H3.3WT cells (LAD-same), whereas 3% (92Mb) underwent LAD loss and 4% (108Mb) underwent LAD gain in at least two of the three H3.3K27M lines (**Fig. 3b**, **Extended Data Fig. 4e**). LAD-same regions exhibited high LaminB1 enrichment, low gene density, and substantial overlap (65%) with majority of LADs identified across a reference panel of twenty H3.3WT cell lines(22,25–27) (**Extended Data Fig. 5a-c)**, consistent with constitutive LADs. In contrast, LAD-loss regions showed reduced overlap with this reference panel and exhibited genomic features characteristic of facultative LADs (**Extended Data Fig. 5a-c**. Thus, H3.3K27M DIPG cell lines recapitulate the widespread and recurrent LAD remodeling observed in primary tumors and that this remodeling preferentially affects facultative rather than constitutive LADs.

### H3.3K27M preferentially localizes to genomic regions that lose lamina association

We next investigated the relationship between H3.3K27M deposition and altered LAD organization. To map the genomic distribution of the H3.3K27M, we performed CUT&RUN using a mutation-specific antibody (anti-K27M) in three H3.3K27M DIPG cell lines (SU-DIPG-XIII, SU-DIPG-XVII, and SU-DIPG-XXIV). All three lines exhibited widespread deposition of the mutant histone across the genome, whereas no signal was detected in H3.3WT control lines (VUMC-DIPG10 and pcGBM2), confirming antibody specificity (**Extended Data Fig. 6a)**.

H3.3K27M was globally depleted in LADs and enriched in non-LAD regions, consistent with preferential deposition in transcriptionally active genomic regions(28) (**Extended Data Fig. 6b**). Notably, among regions that underwent LAD remodeling, LAD-loss regions exhibited strong enrichment of H3.3K27M, whereas LAD-gain and LAD-same regions showed little or no detectable signal (**Fig. 3c**). These findings reveal a close association between local H3.3K27M deposition and loss of lamina association.

We next examined the chromatin states associated with LAD remodeling. As expected, H3K27me3 was globally depleted across all three H3.3K27M cell lines (**Extended Data Fig 6c**), with comparable reductions observed in LAD-loss, LAD-gain and LAD-same regions (**Fig.3c**). In contrast, the active chromatin mark H3K27ac, which is elevated in DIPG(9), was selectively enriched within LAD-loss regions (**Fig. 3c**). Together, these results indicate that regions that detach from the nuclear lamina are distinguished by local H3.3K27M enrichment and acquisition of a more permissive chromatin state characterized by increased H3K27ac.

### LAD loss is more strongly associated with gene activation than H3K27me3 depletion in H3.3K27M DIPG cell lines

To determine how alterations in LAD architecture influence gene expression in DIPG cell lines, we performed RNA-seq on three H3.3K27M-mutant cell lines and two H3.3WT controls, each in biological duplicate. PCA confirmed clear separation by cell line (**Extended Data Fig. 6d**). Consistent with the repressive nature of the nuclear lamina, genes located within LADs were expressed at significantly lower levels than genes residing in non-LAD regions across both H3.3K27M and H3.3WT cells (**Extended Fig. Data Fig. 6e**).

Genes located within LAD-loss regions exhibited robust transcriptional upregulation in H3.3K27M-mutant cells compared to H3.3WT controls (**Fig. 4a; Extended** ). In contrast, genes within regions of H3K27me3 loss showed comparatively modest increases in expression (**Fig. 4b**). GO analysis of upregulated genes within LAD-loss regions revealed enrichment for neuronal programs including, neuronal projection guidance, dendrite morphogenesis, and postsynapse organization (**Extended Data Fig. 6f**).Representative LAD-loss loci, including DIPG associated genes *PTPRZ1* and *PTN*, neuronal genes *ADGRL3, ASTN1*, and *LSAMP,* and proliferation-associated genes *CDK6* and *EGFR* showed loss of lamina association accompanied by increased H3.3K27M incorporation, elevated H3K27ac and strong transcriptional upregulation **(Fig. 4c, Extended Data Fig. 6g).** Together, these findings indicate that loss of lamina association preferentially activates neuronal and cell cycle gene programs and is more strongly associated with transcriptional activation than H3K27me3 depletion in H3.3K27M DIPG cells.

**Figure 4.**
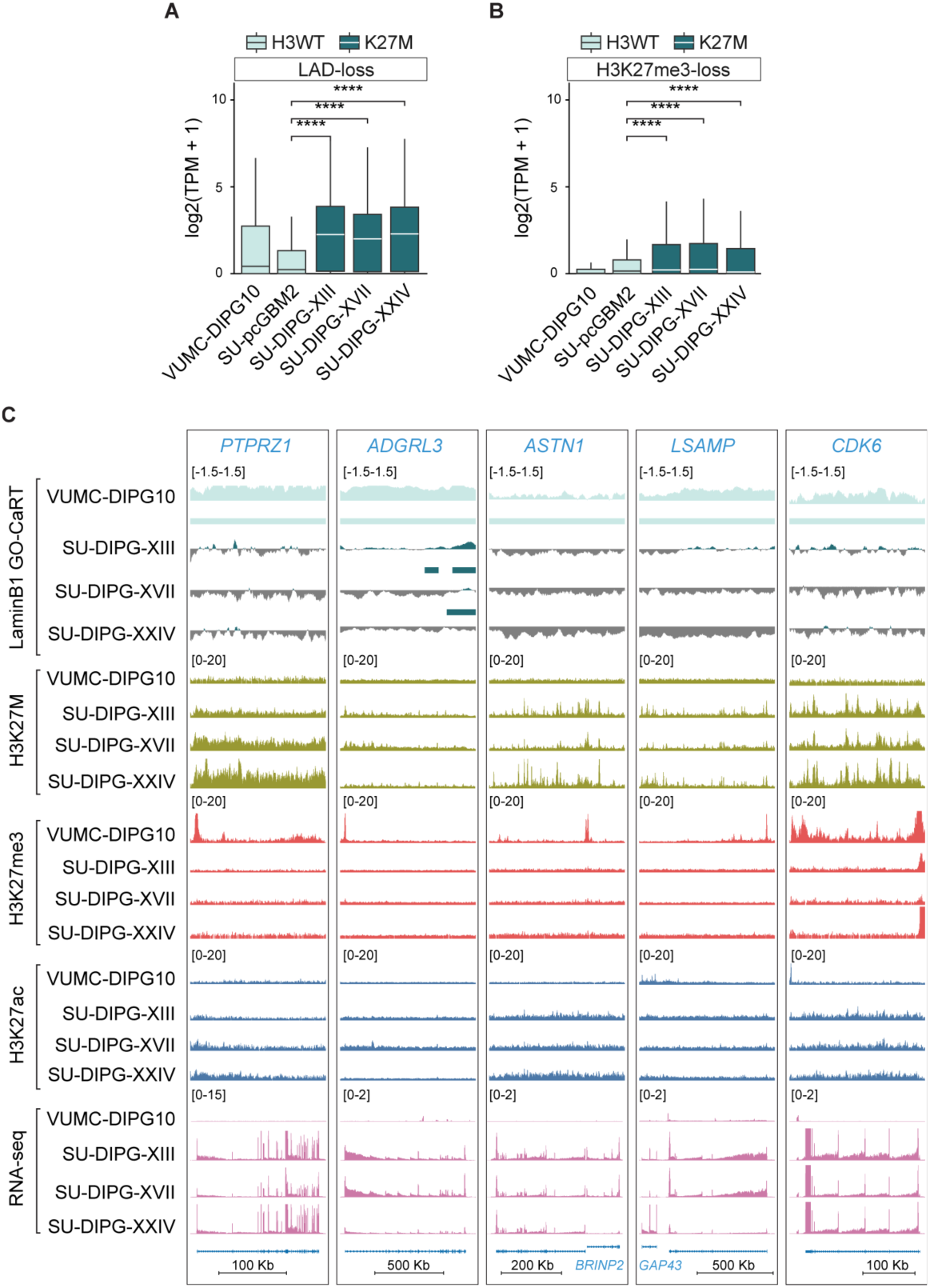
LAD loss leads to larger absolute gene upregulation compared to H3K27me3 loss **A, B,** Boxplots of gene expression represented as log_2_ (TPM + 1) for genes in consensus LAD-loss regions (**A)** or H3K27me3-loss regions **(B**) in VUMC-DIPG10, SU-pcGBM2, SU-DIPG-XIII, SU-DIPG-XVII, and SU-DIPG-XXIV. Consensus loss is defined as loss in at least 2 of 3 H3K27M-positive DIPG cell lines relative to both H3WT controls. Significance was assessed using unpaired Wilcoxon rank-sum test comparing each H3K27M cell line with SU-pcGBM2 as indicated by brackets. ****, P ≤ 0.0001. **C** Representative genome browser tracks of consensus LAD loss regions containing *PTPRZ1, ADGRL3, ASTN1, LSAMP, and CDK6*. Top tracks show log2(LMNB1/IgG) signal per cell line. H3K27M, H3K27me3, and H3K27ac CUT&RUN tracks are shown as normalized reads per genomic content for each cell line. RNA-seq signal is shown in bins per million.

### H3.3K27M expression is sufficient to remodel LADs in human stem cell models

To determine whether H3.3K27M is sufficient to alter LAD organization, we engineered H1 embryonic stem cells (ESCs) with piggyBac-integrated transgenes encoding HA-tagged wild-type H3.3 or H3.3K27M. Engineered ESCs were differentiated into neural precursor cells (NPCs), followed by transgene induction, and profiling by LaminB1 GO-CaRT, CUT&RUN, and RNA-seq (**Fig. 5a**). Transgene expression was confirmed by immunostaining and western blotting and NPC identity was verified by expression of established marker genes (**Extended Data Fig. 7 a-c**). CUT&RUN demonstrated robust H3.3K27M incorporation and the expected global reduction in H3K27me3 in H3.3K27M-expressing NPCs (**Fig. 5b, c**).

**Figure 5.**
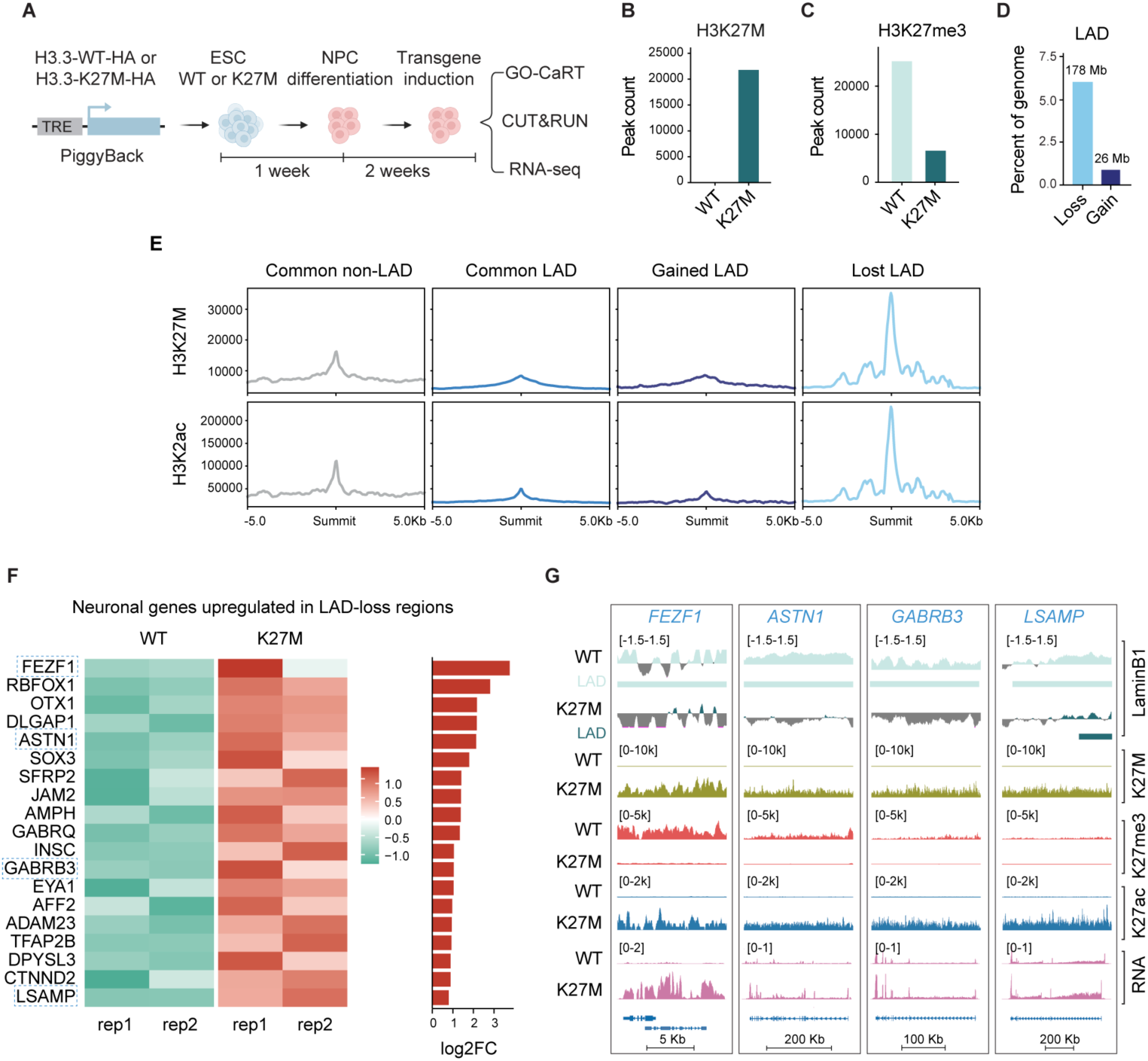
H3.3K27M expression leads to LAD remodeling and neuronal gene activation in transgenic neural precursor cells. **A,** Experimental overview of engineered hESC carrying doxycycline-inducible H3.3K27M-HA or H3.3WT-HA via piggyBac genomic integration. Cells were differentiated into NPCs for one week and H3.3 transgene expression was induced the subsequent two weeks. Cells were harvested for LaminB1 GO-CaRT, CUT&RUN, and RNA-Seq. **B, C,** Number of H3K27M (**B**) or H3K27me3 (**C**) peaks present in H3.3WT NPCs or H3.3K27M NPCs. **D,** Percentage of analyzed genome classified as LAD-loss or LAD-gain in H3.3K27M NPCs relative to H3.3WT NPCs. **E**, Average H3K27M (top row) and H3K27ac (bottom row) signal profiles centered on H3K27ac summits flanked by 5 kb on either side stratified by H3.3K27M NPC LAD changes relative to H3.3WT NPC LADs. **F,** Heatmap of normalized expression of select neuronal genes upregulated in LAD-loss regions with two replicates per genotype. Juxtaposed bar plots show log2 fold change in H3.3K27M NPCs relative to H3.3WT NPCs of corresponding genes. **G,** Representative genome browser views of *FEZF1, ASTN1, GABRB3, and LSAMP.* Signal tracks for LaminB1 are shown as log2(LMNB1/IgG). Signal track for H3K27M, H3K27me3, and H3K27ac are shown as spike-in normalized coverage signal. RNA-seq is presented as bins per million.

LaminB1 GO-CaRT profiling revealed that LADs occupied 46.1% (H3.3WT) and 40.9% (H3.3K27M) of the genome and retained typical heterochromatin features (**Extended Data Fig. 7d, e**). Notably, H.3K27M-NPCs exhibited marked LAD remodeling following transgene induction, with 178 Mb (∼6% of the genome) of LAD-loss and 26 Mb (∼0.9% of the genome) of LAD-gain relative to H3.3WT NPCs (**Fig. 5d**). Thus, H3.3K27M-induced LAD remodeling was strongly biased toward loss of lamina association.

To assess differences in the local incorporation of K27M across LAD-loss, LAD-gain, LAD-same and non-LAD regions, we examined all chromatin regions with maximal enrichment of H3K27ac termed summits. These H3K27ac summits largely corresponded to active enhancer regions and the genomic regions of actively transcribed genes (**Extended Data Fig. 7f**). As expected, in non-LAD regions, K27M levels were higher than in LAD-same regions (**Fig. 5e**). The small fraction of the genome that was LAD-gain had low levels of K27M (**Fig. 5e**). In contrast, in LAD-loss regions, K27M incorporation was very high, exceeding the levels of genes in non-LAD regions, linking local H3.3K27M incorporation to regions that disengage from the nuclear lamina (**Fig. 5e**).

We next examined the transcriptional consequences of LAD remodeling. PCA of RNA-seq separated NPCs according to H3.3 status (**Extended Data Fig. 7g**). As expected, genes within LADs were expressed at significantly lower levels than genes in non-LADs in both H3 WT and H3K27M NPCs (**Extended Data Fig.7h**). Genes within LAD-loss regions showed a modest overall increase in expression in H3.3K27M NPCs (**Extended Data Fig. 7i**). Notably, a subset of neuronal genes within these regions exhibited robust transcriptional upregulation, including *FEZF1, RBFOX1, OTX1, DLGAP1, ASTN1, GABRB3*, and *LSAMP* (**Fig. 5f**). Representative loci, including *FEZF1*, *ASTN1*, *GABRB3*, and *LSAMP*, showed loss of LaminB1 association accompanied by H3.3K27M incorporation, depletion of H3K27me3, increased H3K27ac, and increased transcription (**Fig. 5g)**. In contrast, genes within common non-LAD regions showed even more subtle but statistically significant changes in expression between H3.3WT and H3.3K27M NPCs, despite H3.3K27M incorporation at these regions (**Extended Data Fig. 7j, k)**. Together, these results demonstrate that H3.3K27M expression is sufficient to remodel LADs, predominantly through loss of lamina association and upregulate neuronal gene programs.

## Discussion

The H3.3K27M oncohistone is widely recognized for its ability to inhibit PRC2 activity and globally reduce H3K27me3 levels. However, the mechanisms linking this epigenetic alteration to the transcriptional programs that drive DIPG remain poorly understood. In this study, we identified disruption of nuclear lamina-associated genome organization as a previously unrecognized consequence of H3.3K27M expression. Across primary DIPG specimens, patient-derived cell lines, and engineered NPCs, H3.3K27M was associated with widespread and recurrent remodeling of LADs. Regions that lost lamina association showed substantially greater transcriptional activation than regions defined by H3K27me3 loss alone, linking reorganization of LADs to the transcriptional consequences of H3.3K27M. Together, our findings expand current models of H3.3K27M-driven epigenetic dysregulation by implicating genome–lamina organization as an important layer of oncogenic transcriptional control.

Altered nuclear morphology and heterochromatin organization are longstanding hallmarks of cancer(14), yet studies of H3.3K27M have largely focused on changes in histone modifications(5), enhancer activity(29), and transcription factor networks(4). Our findings extend this framework to higher-order spatial genome organization. Despite relatively stable overall LAD coverage, H3.3K27M cells exhibited extensive redistribution of lamina-associated chromatin. Notably, this remodeling preferentially affected facultative LADs, whereas constitutive LADs remained comparatively stable. Because facultative LADs are developmentally regulated and vary across cell states(18,30), their selective disruption suggests that H3.3K27M preferentially acts on regions of the genome with inherent developmental plasticity. Consistent with this model, LAD-loss regions were enriched for neuronal and developmental genes, suggesting that disruption of peripheral genome organization may facilitate inappropriate engagement of lineage-specific transcriptional programs in DIPG.

A central finding of this study is that loss of lamina association is more closely associated with transcriptional activation than H3K27me3 depletion. Genes within LAD-loss regions were consistently upregulated in primary tumors and patient-derived cell lines, whereas genes within regions defined by H3K27me3 loss showed comparatively modest transcriptional changes. This distinction is particularly important given the near-global depletion of H3K27me3 caused by H3.3K27M. Indeed, previous studies have shown that widespread H3K27me3 loss does not fully account for the transcriptional changes induced by H3K27M(7,8), suggesting that additional regulatory mechanisms determine which genes become activated. Our findings suggest that the spatial context of chromatin provides an additional layer of regulatory information, with release from the nuclear lamina identifying genomic regions particularly permissive to transcriptional activation. This model is consistent with our recent findings demonstrating that nuclear lamina acts as key determinant of transcriptional control independently of H3K27me3(22) and provides a potential explanation for why global H3K27me3 depletion alone incompletely predicts the transcriptional consequences of H3.3K27M.

The identity of genes released from lamina-associated repression may have important implications for DIPG biology. LAD-loss regions were enriched for pathways involved in axonogenesis, synapse assembly, and cell adhesion, and included neuronal genes that became strongly activated following lamina detachment. This is particularly relevant given evidence that DIPG and other gliomas engage neuronal developmental programs and form functional neuron– glioma synapses that promote tumor growth through activity-dependent signaling(23,24,31). Our findings raise the possibility that H3.3K27M-associated LAD remodeling facilitates access to neurodevelopmental and synaptic gene programs that support these tumor–neuronal interactions. More broadly, selective remodeling of facultative LADs may provide a mechanism through which H3.3K27M exploits the developmental plasticity of glioma precursor cells to reinforce transcriptional states favorable to tumor progression.

Our data also suggest that H3.3K27M contributes directly to local disruption of genome– lamina interactions. Although the mutant histone was globally depleted from LADs, it was strongly enriched in regions that underwent LAD loss. The engineered NPC model provides further evidence that LAD remodeling is an intrinsic consequence of H3.3K27M expression rather than solely a secondary feature acquired during tumor evolution. Expression of H3.3K27M was sufficient to induce extensive LAD reorganization, with remodeling strongly biased toward loss of lamina association. These findings support a model in which H3.3K27M acts on developmentally plastic chromatin to reorganize genome–lamina interactions and establish a chromatin environment permissive for aberrant neuronal transcriptional programs.

In summary, our study identifies disruption of nuclear lamina-associated genome organization as a previously unappreciated dimension of H3.3K27M-mediated epigenetic dysregulation. H3.3K27M reshapes the transcriptional landscape not only through global perturbation of histone modifications, but also through selective reorganization of developmentally regulated chromatin at the nuclear periphery. These findings expand current models of DIPG pathogenesis, establishing spatial genome organization as a key mechanism of transcriptional re-programming in H3.3K27M-driven cancers. More broadly, our work provides a framework for understanding how an oncohistone can convert widespread epigenetic disruption into selective oncogenic transcriptional outputs.

## Methods

### Tumor tissue acquisition and processing

Deidentified primary DIPG tumor specimens (SF-5475, SF-5599, and SF-5606) were obtained from the UCSF Brain Tumor Center (BTRC) with informed consent and in accordance with Institutional Review Board (IRB)-approved protocols at the University of California, San Francisco (UCSF). Postmortem brain tumor tissues were dissected into small fragments and snap-frozen immediately at the time of autopsy.

### Cell cultures

Patient-derived H3.3K27M DIPG cell lines (SU-DIPG-XIII, SU-DIPG-XVII and SU-DIPG-XXIV) and H3WT pediatric glioma cell line, pcGBM2 were generously provided by the laboratory of Michelle Monje (Stanford University) and have been previously described(32) . VUMC-DIPG10, an H3WT DIPG cell line, was generously provided by Dr. E(33). All cultures were grown as neurospheres in Tumor Stem Media (TSM) consisting of a 1:1 mixture of DMEM/F12 (Invitrogen) and Neurobasal-A (Invitrogen), supplemented with 1X B27(-A) (Invitrogen), human-bFGF (20 ng/ml) (Shenandoah Biotech), human-EGF (20 ng/ml) (Shenandoah), human PDGF-AB (20 ng/ml) (Shenandoah) and heparin (10 ng/ml). The H1 ESC line was obtained from WiCell (Madison WI). ESCs were cultured on Matrigel-coated 6-well plates in StemFlex medium (Gibco) supplemented with 10 μM Y-27632 ROCK inhibitor (ROCKi) during the first 16–20 h after plating. Cells were fed fresh medium daily until reaching 80–90% confluency, at which point they were passaged as cell aggregates using ReLeSR (STEMCELL Technologies).

### Generation of H3.3K27M or H3.3 wildtype stable ESC lines

The PiggyBac vector, pB-TRE-EGFP-EF1a-rtTA (addgene #104454) was digested with *NheI* and *AgeI* to remove the EGFP cassette and linearize the plasmid backbone. cDNAs encoding wild-type H3F3A (H3.3WT) and mutant H3F3A-K27M, each containing a C-terminal 3xHA tag, were PCR-amplified from synthetic gene blocks (Integrated DNA Technologies, IDT) and cloned into the linearized vector using the In-Fusion HD Cloning Kit (Takara Bio). The resulting constructs, pB-TRE-H3F3A-WT-3xHA and pB-TRE-H3F3A-K27M-3xHA, also contained a constitutive EF1α promoter driving puromycin resistance for selection.

ESCs were transfected using the Neon Transfection System (Thermo Fisher Scientific). Briefly, cells were dissociated into single cells using Accutase, and 5 x 10^5^ cells were used per transfection. Cells were washed with PBS, centrifuged at 300 x *g* for 3 min, and resuspended in 10 μL of Neon Resuspension Buffer (R Buffer). For each transfection, 1.5 μg of PiggyBac expression construct and 0.5 μg of PiggyBac transposase plasmid (3:1 molar ratio) were mixed in a PCR tube. Cell mixture prepared in R Buffer (10 μl) were added to the DNA mixture and gently mixed. Electroporation was performed using a 10 μL Neon tip with the following settings: Voltage, 1400V; Width, 20 ms; Pulses, 1. Using the Neon tip, electroporated cells were transferred into pre-warmed Matrigel-coated 6-well plates containing StemFlex medium supplemented with ROCK inhibitor. The following day, the medium was replaced with fresh StemFlex medium containing ROCK inhibitor. Puromycin selection (1 μg/mL) was initiated 72 h after electroporation and maintained for one week. Following selection, 10,000 ESCs were plated onto 10-cm dishes, and colonies derived from single cells were picked under a phase-contrast microscope and expanded as clonal cell lines. Expression of H3.3WT or H3.3K27M was induced by the addition of doxycycline (1 μg/mL) to the culture medium.

### Differentiation of ESCs to NPCs

H3.3WT and H3.3K27M ESCs were differentiated into NPCs using the StemXVivo Neural differentiation kit (R&D systems) according to the manufacturer’s instructions. Briefly, ESCs were dissociated using ReLeSR and plated as small aggregates onto a Matrigel-coated 6-well plate in StemFlex media containing 10 μM ROCKi. The following day, the media was replaced with NPC differentiation Media (Day 0). Cells were fed fresh NPC differentiation media for seven days. On Day 8, NPCs were dissociated and replated in StemFlex media containing 10 μM ROCKi. The next day, doxycycline (1μg/mL) was added to the culture media to induce the expression of H3.3WT or H3.3K27M. Induction was maintained for two weeks, with doxycycline-containing medium replaced every other day.

### Nuclei isolation from DIPG tumor specimens and FANS

Nuclei isolation and FANS were performed as described previously(22). Snap-frozen postmortem DIPG tumor specimens were cut into small pieces and transferred to a pre-chilled 7-mL Dounce Tissue Grinder (Wheaton) containing 5 mL nuclei extraction buffer (NEB: 10 mM HEPES pH 7.4, 25 mM KCl, 5 mM MgCl_2_, 0.25 M sucrose, 0.1% Triton X-100, 1x Halt protease inhibitor cocktail (Thermo Fisher)). RiboLock (Thermo Fisher) was added to all the buffers to preserve RNA integrity. Samples were homogenized on ice with 5-6 strokes using loose pestle A followed by 8-10 strokes using tight pestle B, until no tissue pieces were visible. The homogenate was incubated on ice for 5 min and then transferred to a pre-chilled 15 mL conical tube. Nuclei were pelleted by centrifugation at 500 *g* for 10 min at 4°C. The supernatant was discarded, and the pellet was resuspended in 10 mL of NEB without Triton X-100. The suspension was filtered through a 40 μm cell strainer into a 50 mL conical tube. Formaldehyde was added to a final concentration of 0.1%, and nuclei were fixed for 2 min at room temperature with gentle rotation. The reaction was quenched by adding glycine to a final concentration of 75 mM. BSA was added to a final concentration of 1%, followed by centrifugation at 500 *g* for 10 min at 4 °C. The supernatant was discarded, and the nuclear pellet was resuspended in 1 mL staining buffer (PBS containing 1% BSA). The suspension was filtered through a 40 μm strainer into a low-bind 1.5 mL microcentrifuge tube, and nuclei were counted under the microscope. Nuclei were pelleted at 500 *g* for 10 min at 4 °C and resuspended in 100-150 μL of staining buffer. Samples were stained at 4 °C on rotation for 1 hr with H3K27M-AlexaFluor 488 antibody (Cell Signaling Technologies, #85023S). Following staining, 900 μL of staining buffer was added and the samples were centrifuged at 500 *g* for 10 min at 4 °C. The nuclear pellet was resuspended in 1 mL of staining buffer and centrifuged at 500 *g* for 10 min at 4 °C. The final pellet was resuspended in 1-2 mL of staining buffer (depending upon the starting material and nuclei yield) and filtered into a 70 µm mesh FACS tube (BD). DAPI was added at 1 μg/ml just before sorting. FANS was performed on BD FACS Aria II sorter using a 70 μm nozzle. Sorted nuclei were collected in 5 mL tubes containing 300-500 μL of collection buffer (PBS containing 5% BSA and RNasin Plus RNase inhibitor (Promega)). Following sorting, nuclei were collected by centrifuging at 500 *g* for 10 min at 4 °C and processed for downstream experiments (RNA-seq, Fab blocking and GO-CaRT/CUT&RUN).

### Fab blocking, GO-CaRT and CUT&RUN analyses

For nuclei isolated by FANS, samples were first subjected to Fab blocking as previously described(22), before proceeding with GO-CaRT and CUT&RUN analyses. Briefly, sorted nuclei were resuspended in PBS containing 1% BSA and aliquoted into 100 μL volumes in 0.5 mL PCR tubes. Activated BioMagPlus Concanavalin A beads (8 μL; Polysciences) were added to each sample, and nuclei were allowed to bind for 10 min at room temperature with rotation. Samples were then placed on a magnetic stand, and the supernatant was removed. To block antibodies used in FANS, nuclei bound to concanavalin A beads were resuspended in 50 μL wash buffer (20 mM HEPES-KOH pH 7.5, 150 mM NaCl, 0.1% BSA, 0.5 mM spermidine and 1× Halt protease inhibitor cocktail) containing anti-rabbit monovalent Fab fragments (Jackson Immuno) at 1:30 dilution. Samples were incubated on a nutator at 4 °C for 30 min. Samples were placed on a magnetic stand, the supernatant was removed, and beads were washed once with 200 μL wash buffer to remove excess Fab fragments.

For all other samples, including cultured cell lines, nuclei were isolated as previously described(20) and directly bound to activated BioMagPlus concanavalin A beads for 10 min at room temperature. All subsequent steps were performed identically for FANS-isolated and cultured-cell nuclei.

Bead-bound nuclei were resuspended in 50 μL antibody-binding buffer (wash buffer containing 2 mM EDTA) containing the primary antibody of interest for GO-CaRT or CUT&RUN. Samples were incubated overnight at 4°C on a nutator. The following day, samples were briefly centrifuged, placed on a magnetic stand, and washed twice with 200 μL wash buffer. Beads were then resuspended in 50 μL wash buffer containing pA/G-MNase and rotated for 1 hour at 4 °C. Samples were briefly spun and placed on a magnetic stand to remove the supernatant. Samples were washed two times with 200 μL wash buffer. Beads were gently resuspended in a 100 μL wash buffer containing 2 mM CaCl_2_ (to activate pA/G-MNase) and placed in a pre-chilled metal block on ice. Digestion was carried out for 30 min and stopped by adding 100 μL of 2XSTOP (200 mM NaCl, 20 mM EDTA, 4 mM EGTA, 50 μg/ml RNase A, 40 μg/ml glycogen). Samples were incubated at 37 °C for 20 min to release pA/G-MNase cleaved fragments. 2 μL SDS (10%) and 2 μL Proteinase K (20 mg/ml) was added to each sample and incubated at 55 °C for 1 hour. DNA was extracted by the phenol-chloroform method. Purified DNA fragments were analyzed by TapeStation High Sensitivity D1000 assay (Agilent).

### Library preparation and sequencing

Sequencing libraries for GO-CaRT and CUT&RUN samples were prepared using KAPA HyperPrep Kit (Roche) following the manufacturer’s instructions. The libraries were amplified for 12–14 PCR cycles. Library quality and fragment size distributions were assessed using the TapeStation D1000 assay (Agilent), and DNA concentrations were quantified using the Qubit High Sensitivity dsDNA assay (Invitrogen). Libraries were pooled and sequenced using 150-bp paired-end reads on either the NovaSeq 6000 or NovaSeq X Plus platform (Illumina).

### GO-CaRT and CUT&RUN alignment

For GO-CaRT and CUT&RUN experiments, paired end reads were first trimmed with bbduk (BBMap v39.1) against BBMap adapter set BBMap adapter reference, using right-end k-mer trimming with *ktrim=r k=23 mink=11 hdist=1 tpe tbo* (*34*). Trimmed reads were then aligned to hg38 (Gencode 39) using bowtie2 (v2.5.4) (35) and the following parameters: *--no-unal --local -- very-sensitive-local --no-mixed --no-discordant \ -q --phred33 -I 10 -X 700.* Reads were then sorted by name using samtools (v1.16.1)(36) collate, mate score fixing with samtools fixmate. Blacklisted regions from (37) were removed using samtools view and duplicates were removed using samtools sort for coordinate sorting followed by markdup.

### Bigwig Processing

#### Coverage Signal: Spike In Normalized

Read coverage files were generated using deeptools bamCoverage v3.5.5(38) with the following parameters: *--normalizeUsing None --extendReads –exactScaling --skipNAs – maxFragmentLength 700 –scaleFactor {factor}*. The scale factor is 10,000 divided by the number of reads aligned to e coli genome from that sample(39)Trimmed reads were aligned to U00096.3 (UCSC) with the same parameters used for alignment to human genome.

#### Coverage Signal: RPGC Normalized

Read coverage files were generated using deeptools bamCoverage with the following parameters: *--skipNAs --normalizeUsing RPGC --effectiveGenomeSize {efgs} --extendReads – exactScaling*. EffectiveGenomeSize was defined per deeptools recommendation for hg38.

#### Log 2 Fold Change Signal

Log 2 Fold Change bigwigs were generated using deeptools bamCompare with the following parameters: *--operation log2 --exactScaling --extendReads --maxFragmentLength 700* data processing and domain calling

### LAD Calling

LADs were called with custom gaussian hmm model identifying Inter-LAD and LAD states inspired by(27). Fixed seeds were used for all random number generation. Briefly, the genome was binned to 10kb and scored using deepTools bamCompare to calculate log2 foldchange for LMNB1 over IgG per replicate with parameters described above with the addition of – *skipZeroOverZero.* Individual log2 fold changes were then z-scored and any bins not part of at least a 100 kb contiguous run with valid scores for all replicates and not blacklisted were removed. Emission and transition parameters were initialized by k-means on a random genome-wide subsample and fitted via Baum–Welch EM with a weak Dirichlet prior on the transition matrix (α_T_ =1.1) and variance shrinkage toward the global variance (λ=0.2) to prevent degenerate state collapse. State paths were assigned by Viterbi decoding. The two states were ordered by mean emission, labelled Inter-LMNB1 (low) and LMNB1 (high), and adjacent bins of identical state merged into domains.

#### Differential LADs

For primary tumor nuclei LAD-loss and LAD-gain were calculated using plyranges setdiff_ranges. For patient derived cell lines consensus LAD-loss regions were defined as LADs in both pcGBM2 and VUMC-DIPG10 but not LAD in at least 2 of 3 H3K27M DIPG patient derived cell lines. LAD-gain was defined analogously. For engineered NPCs, LAD-loss and LAD-gain were calculated in the same manner as primary tumor nuclei.

### Domain and peak calling

H3K27me3 peaks were calculated with epic2(40) using bam files as input and the following parameters: *--guess-bampe -a -gn hg38 -d ’GL*|KI*|JH*|chrM|MT’ -bin 2500 -g 3 -fdr 0.05*

H3K27M, H3K27ac, and H3K4me3 were all calculated with macs2(41) callpeak function with bam files as input and the following parameters: *-f BAMPE -g 2.9e9 --seed 100.* For both epic2 and macs2 CUT&RUN samples against IgG served as control samples and all replicates were used in peak calls.

### RNA isolation and RNA-seq

Total RNA was extracted from sorted nuclei using the Quick-RNA FFPE RNA Extraction Kit (Zymo Research). Total RNA from patient-derived cell lines and NPCs was extracted using the Direct-zol RNA Miniprep Kit (Zymo Research). Ribosomal RNA (rRNA) was depleted using the NEBNext rRNA Depletion Kit v2 (New England Biolabs, NEB). RNA-seq libraries were prepared using the NEBNext Ultra II Directional RNA Library Prep Kit (NEB) following the manufacturer’s instructions. Library quality and fragment size distributions were assessed using the TapeStation D1000 assay (Agilent), and DNA concentrations were quantified using the Qubit High Sensitivity dsDNA assay (Invitrogen). Libraries were pooled and sequenced using 150-bp paired-end reads on either the NovaSeq 6000 or NovaSeq X Plus platform (Illumina).

### Gene expression analyses

Paired end reads were trimmed using BBduk settings described above. Transcript abundance was quantified using salmon (v1.10.1) (42) quant with selective alignment index generated from Gencode v39 annotations(43). The following parameters were used in salmon quant: *--mimicBT2 --numBootstraps 100 --seqBias –gcBias*. Downstream gene expression analysis was performed in R (v4.3.3) with packages tximeta (v1.20.3)(44) and DESeq2 (v1.42.1)(45).

### Gene ontology analyses

Gene ontology analyses were performed in R with the function enrichGO from the module clusterProfiler (v4.10.1)(46) to identify enriched biological processes from list of ensembl gene ids with entire genome set as background. Only GO terms with qvalue < 0.05 were deemed statistically significant.

### Statistics and data reproducibility

All GO-CaRT and RNA-seq experiments in patient derived cell lines and ESC derived precursors were performed with at least 2 biological replicates. No statistical test was used to predetermine the sample sized used. P values were derived from two-sided Wilcoxon’s ran sum test for comparisons throughout the study.

### Immunofluorescence

H3.3K27M and H3.3WT H1-hESCs were cultured on Matrigel-coated chambered slides and the expression of transgene was induced with doxycycline (1μg/mL) for 3 days. Cells were fixed with 4% paraformaldehyde for 20 min at RT, followed by quenching with 125 mM glycine. Cells were then permeabilized and blocked (5% normal donkey serum, 5% normal goat serum [Jackson Immunoresearch], 0.25% Triton X-100 in PBS) for 1 h at RT. Cells were incubated with rabbit anti-HA (Abcam #ab16048, 1:500) or rabbit anti-H3K27M (Abcam #ab13970, 1:500) primary antibodies, diluted in antibody buffer (5% normal donkey serum, 5% normal goat serum, 0.1% Triton X-100 in PBS) overnight at 4° C. The next day, cells were washed 3x for 10 min in PBST followed by incubation in antibody buffer with secondary antibodies, goat anti-rabbit Alexa Fluor 488 (Invitrogen, all 1:500) for 2 hours at RT in the dark. Cells were then washed 3x in PBST, incubated with 1µg/mL DAPI in PBS for 5 min, and mounted with ProLong Glass Antifade Mountant. Slides were imaged on a Leica Stellaris confocal microscope with 63x objective.

### Western blot

H3.3 WT and H3.3K27M NPCs were washed with ice-cold PBS and harvested on ice in RIPA buffer (50 mM Tris-HCl pH 7.5, 150 mM NaCl, 1 mM EDTA, 0.1% SDS, 0.5% sodium deoxycholate) supplemented with 1X Halt Protease Inhibitor Cocktail. Lysates were sonicated on ice and centrifuged at 3,000 g for 10 min at 4 °C. NuPAGE reducing agent was added to supernatants at a concentration of 10%. Samples were mixed with LDS Sample Buffer (Thermo Fisher) and boiled for 5 min. 20 μg of protein was resolved by SDS-PAGE. Protein was transferred onto PVDF membranes using the iBlot Dry Blotting System, which were then blocked in 5% bovine serum albumin in PBS with 0.1% Tween-20 (PBST) followed by overnight incubation at 4 °C in primary antibodies diluted in blocking buffer: anti-Histone H3-K27M (1:1,000; Abcam, ab190631), anti-HA tag (1:1,000; Abcam, ab9110), and anti-GAPDH (1:5,000; Proteintech, 60004-1-Ig). Membranes were washed in PBST and incubated in HRP-tagged anti-rabbit secondary antibody (Cell Signaling, diluted 1:5000 in blocking buffer) or HRP-tagged anti-mouse secondary antibody (Cell Signaling, diluted 1:5000 in blocking buffer) for one hour at room temperature. Proteins were visualized using SuperSignal West Pico Plus Chemiluminescent Substrate (Thermo Fisher) and imaged on a LICOR Odyssey XF system.

## Data availability

Next generation sequencing data from primary tumor samples will be uploaded to the appropriate controlled access public repository. Next generation sequencing data from patient derived cell lines and ESC derived NPCs will be uploaded to GEO and available upon publication of the manuscript.

## Code availability

Code used for analysis in this manuscript will be published on a public code repository upon publication of the manuscript.

## Extended Data Figures

**Extended Data Figure 1.**
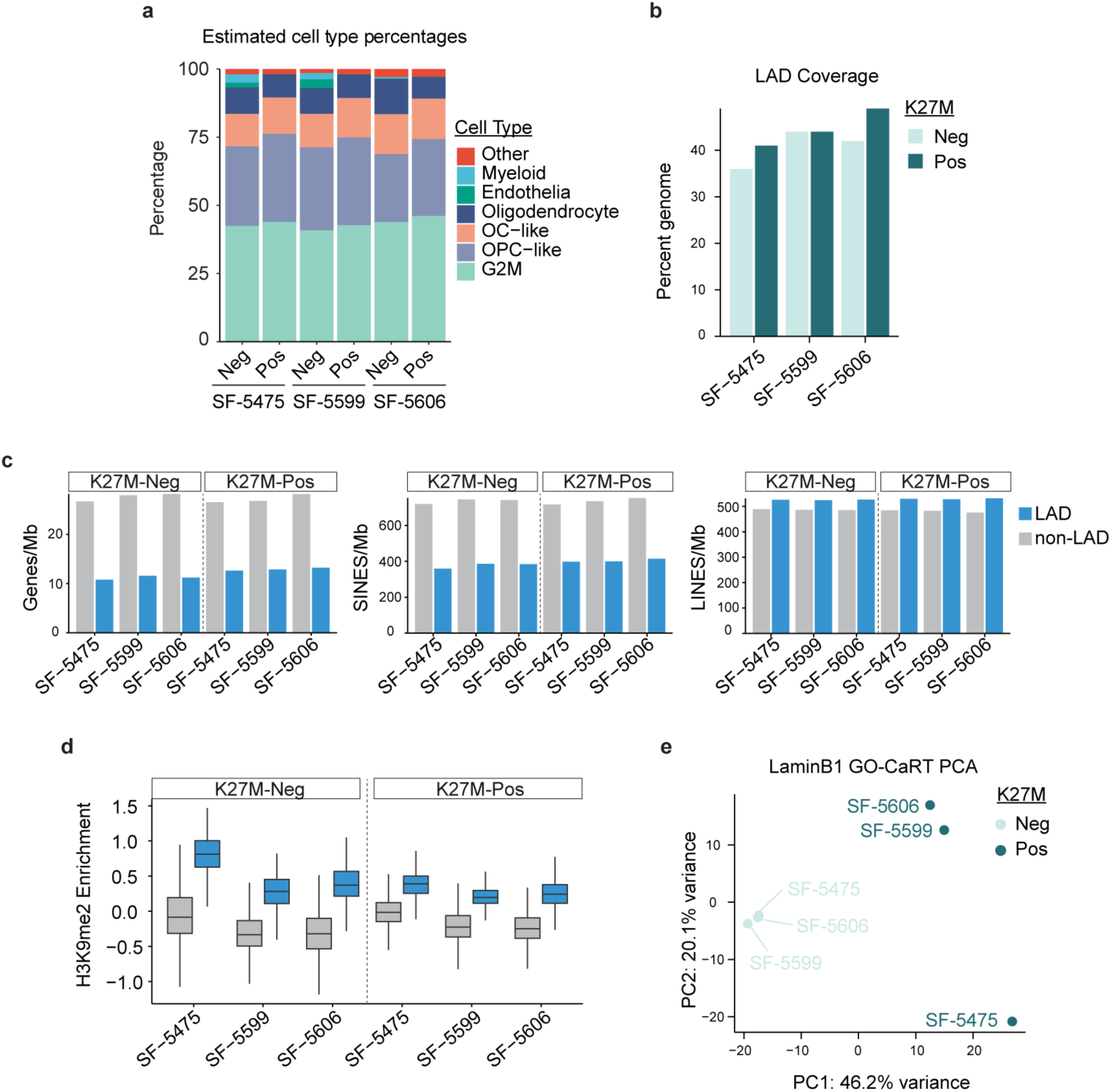
LADs in Primary DIPG Tumor Samples exhibit canonical features. A, Estimated cell type proportions of H3K27M-Negative and H3K27M-positive nuclei from primary DIPG tumors. Proportions were estimated from bulk RNA-seq using MuSiC(47) to deconvolute cell states utilizing pediatric glioma scRNA-Seq dataset as reference. **B,** Percentage of analyzed genome classified as LAD in each tumor-genotype combination. **C,** Gene density, SINE density, and LINE density in LADs and Inter-LADs of each tumor-genotype combination as counts per Mb. **D,** Mean H3K9me2 signal, log2(H3K9me2/IgG), for each tumor faceted by genotype in Inter-LADs (grey) and LADs (blue). **E,** PCA of normalized log2(LMNB1/IgG) signal binned at 100 kb for each tumor-genotype combination. Signal was averaged per replicate prior to quantile normalization. Percentages show percent of variance explained by each principal component.

**Extended Data Figure 2.**
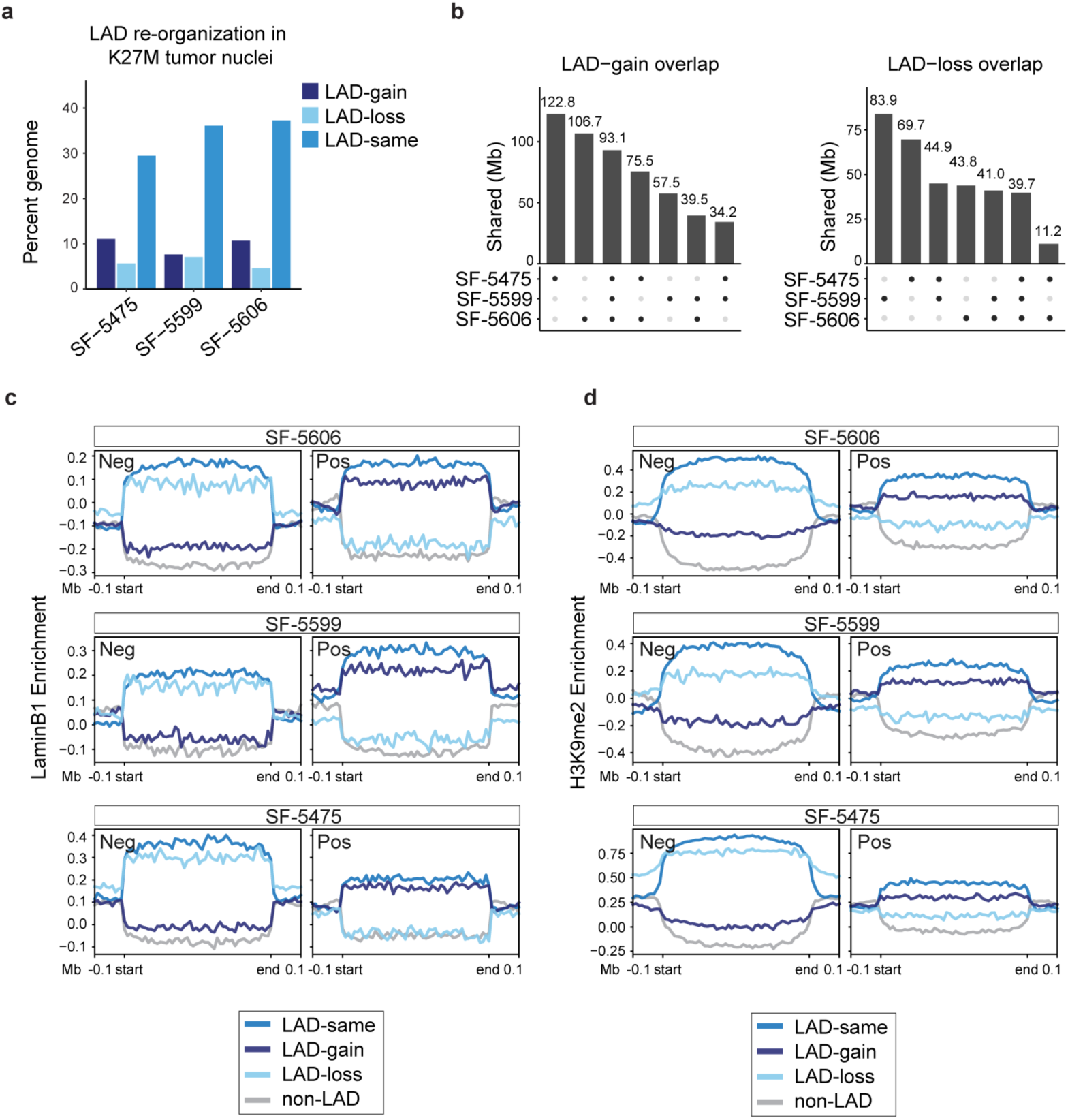
LAD remodeling characteristics in primary tumors. A, Percentage of analyzed genome classified as LAD-gain, LAD-loss, or LAD-same in H3K27M-positive nuclei relative to paired H3K27M-negative nuclei. **B,** UpSet plots showing overlap in Mb of LAD-gain (left) and LAD-loss (right) across all combinations of tumor samples. **C, D,** Average Lamin B1 GO-CaRT (C) and H3K9me2 CUT&RUN signal profiles for H3K27M-positive LAD transitions. Signal is defined as (log2(signal/IgG). Regions are scaled to 500 kb and are flanked by 100 kb regions. Profiles are shown for H3K27M-negative (left) and H3K27M-positive (right) samples.

**Extended Data Figure 3.**
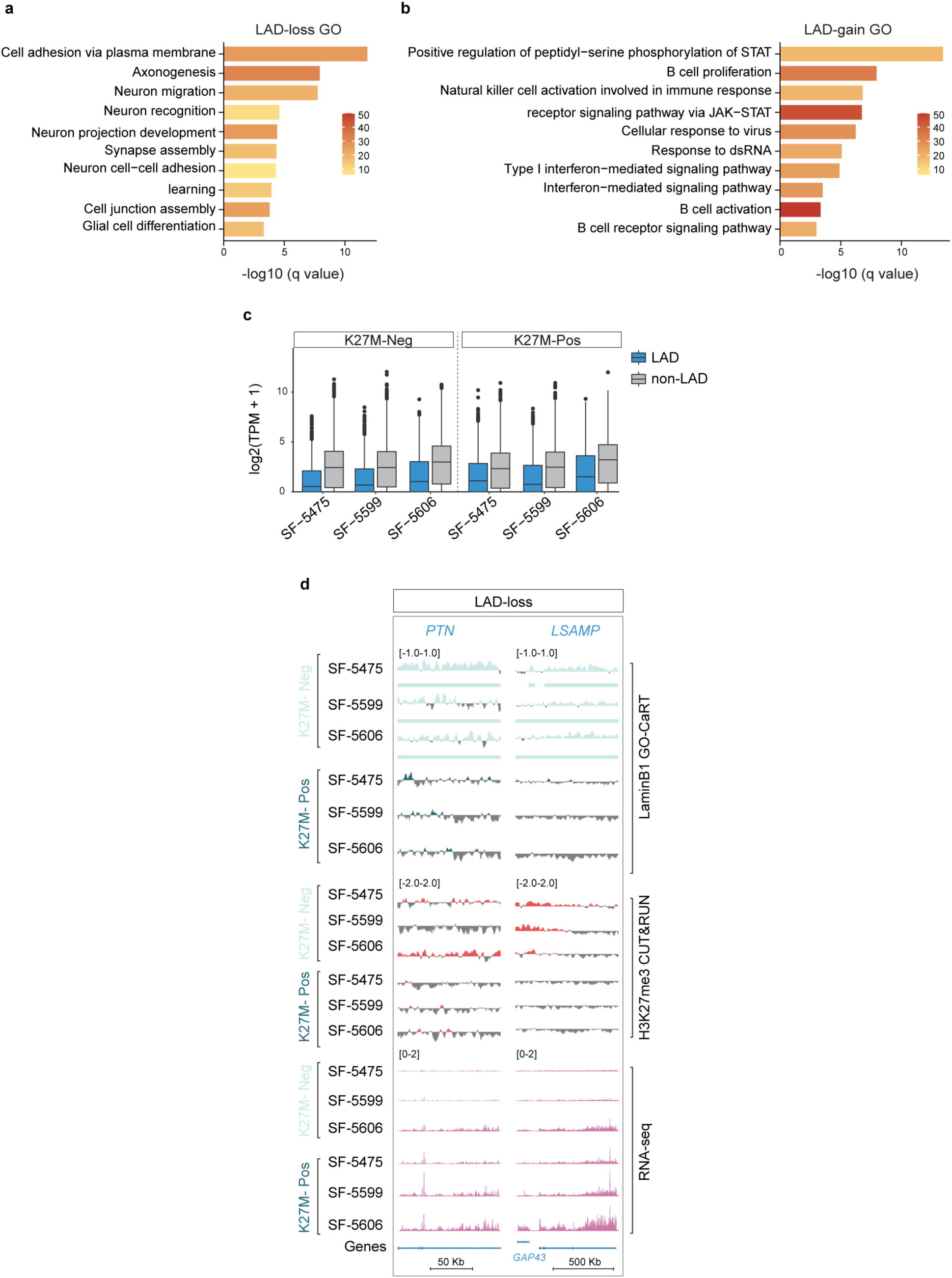
Neuronal genes are enriched and expressed in LAD-loss regions of primary DIPG samples. A, B, Gene Ontology of biological process enrichment for genes in LAD-loss regions (lost in ≥2 of 3 samples) (**A**) and LAD-gain regions (gained in ≥ 2 of 3 samples) (**B**). Color intensity represents number of genes associated with each term. **C,** Boxplots of gene expression, shown as log2(TPM + 1) for genes in LAD and Inter-LAD regions for each tumor faceted by genotype. **D,** Representative genome browser tracks of LAD-loss loci *PTN* and *LSAMP*. Tracks are ordered with H3K27M-negative samples above and H3K27M-positive samples below for each assay. LaminB1 GO-CaRT and H3K27me3 signal is shown as log2(signal/IgG). RNA-seq tracks are shown as bins per million.

**Extended Data Figure 4.**
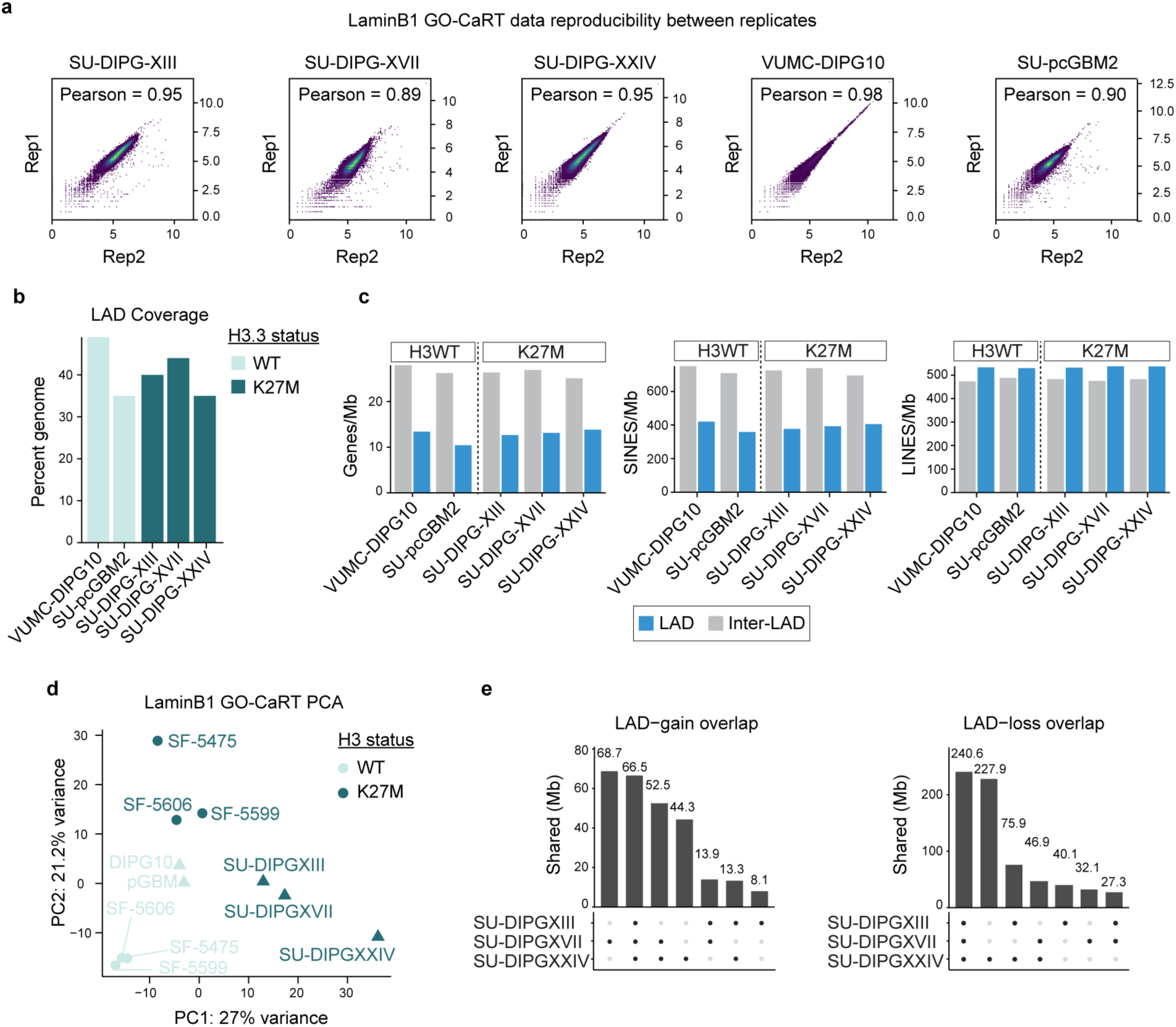
LADs in patient derived DIPG cell lines are similar across replicates and cell lines. A, Spearman correlation of LMNB1 coverage signal between two biological replicates of each patient derived cell line. **B,** Percentage of the analyzed genome classified as LADs in each cell line. **C,** Gene density, SINE density, and LINE density in LADs and Inter-LADs for each cell line as counts per Mb. **D,** PCA of normalized log2(LMNB1/IgG) signal binned at 100 kb for primary tumor samples and patient-derived cell lines. Signals were averaged per replicate prior to quantile normalization. Percentages show percent of variance explained by each principal component. **E,** UpSet plots showing overlap in Mb of LAD-gain (left) and LAD-loss (right) loci from H3K27M DIPG cell lines relative to VUMC-DIPG10.

**Extended Data Figure 5.**
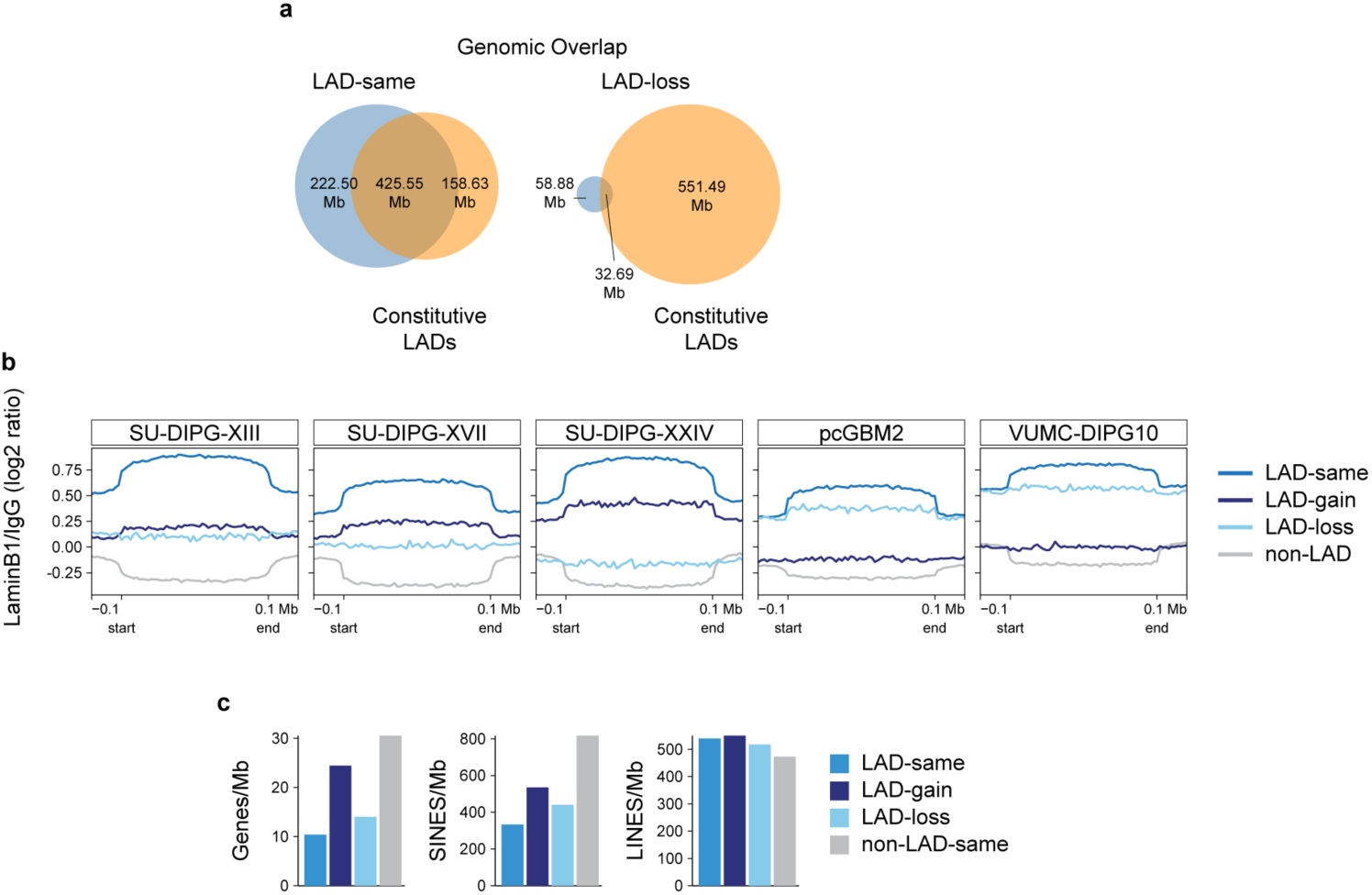
LAD loss regions have facultative LAD characteristics. A, Overlap in Mb of consensus LAD-same (left) and LAD-loss (right) with constitutive LAD reference set, which is derived from LADs of twenty cell lines. **B,** Average LMNB1 GO-CaRT signal in log2(LMNB1/IgG) across consensus LAD-same, LAD-gain, LAD-loss, and common non-LAD regions in H3K27M DIPG patient derived cell lines relative to WT controls. **C,** Gene, SINE, and LINE densities for each LAD transition in counts per Mb.

**Extended Data Figure 6.**
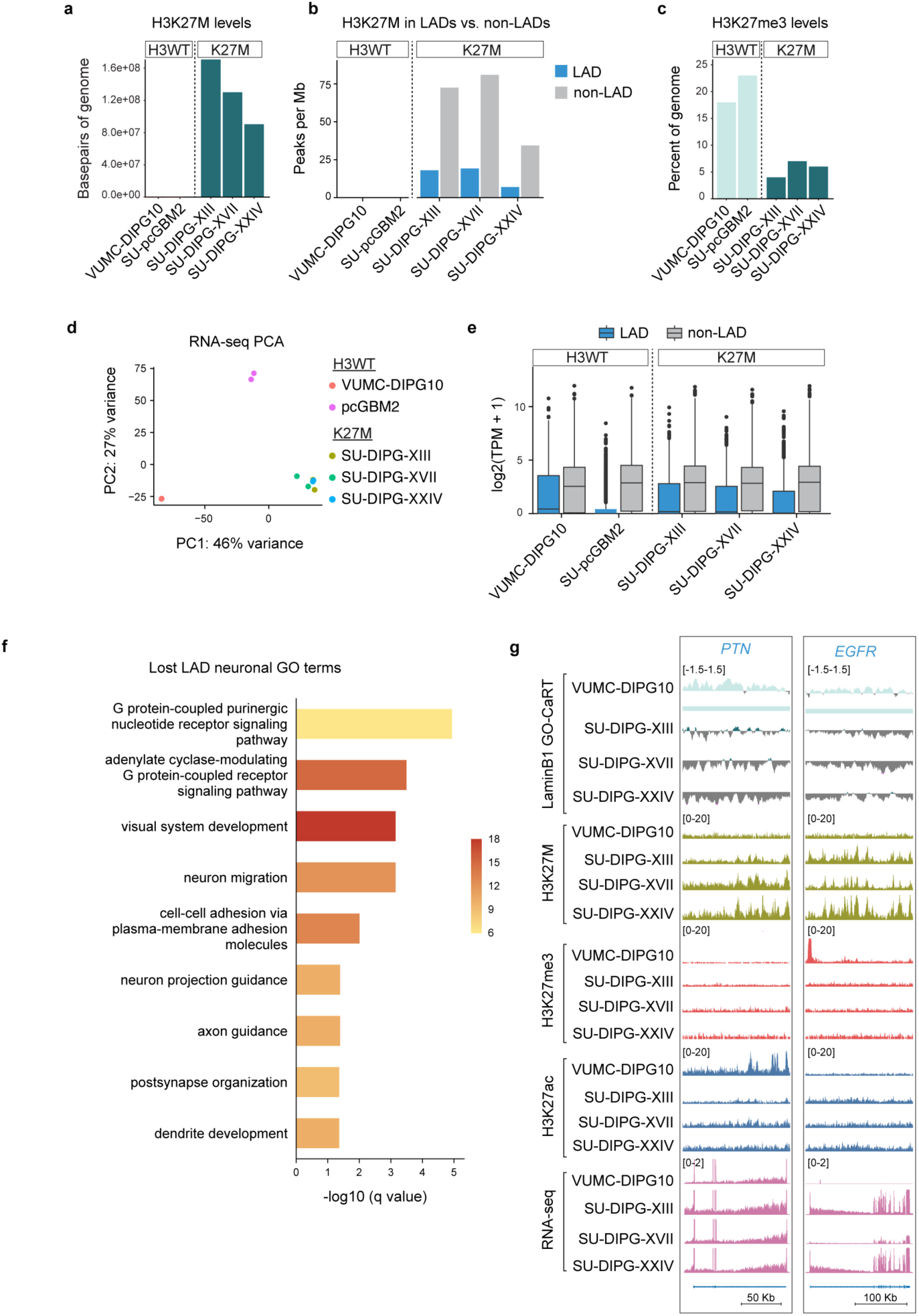
H3K27M expression is associated with upregulation of neuronal genes in LAD-loss regions of patient derived cell lines. A, Genomic coverage in bp of H3K27M peaks in patient derived cell lines. **B,** Peak density of H3K27M in mutant cell lines and WT cell lines shown as peaks per Mb. **C,** Percentage of analyzed genome covered by H3K27me3 peaks per cell line. **D,** PCA of RNA-seq profiles of cell line biological replicates. **E,** Boxplots of gene expression shown as log2(TPM+1) of LAD and non-LAD genes per cell line. **F,** GO of biological processes pertaining to neuronal function in consensus lost LADs of H3K27M DIPG patient derived cell lines. **G,** Representative genome browser tracks of LAD-loss loci *PTN* and *EGFR*. LaminB1 GO-CaRT shows log2(LMNB1/IgG). H3K27M, H3K27me3, and H3K27ac CUT&RUN tracks show normalized coverage as RPGC. RNA-seq tracks show coverage in bins per million.

**Extended Data Figure 7.**
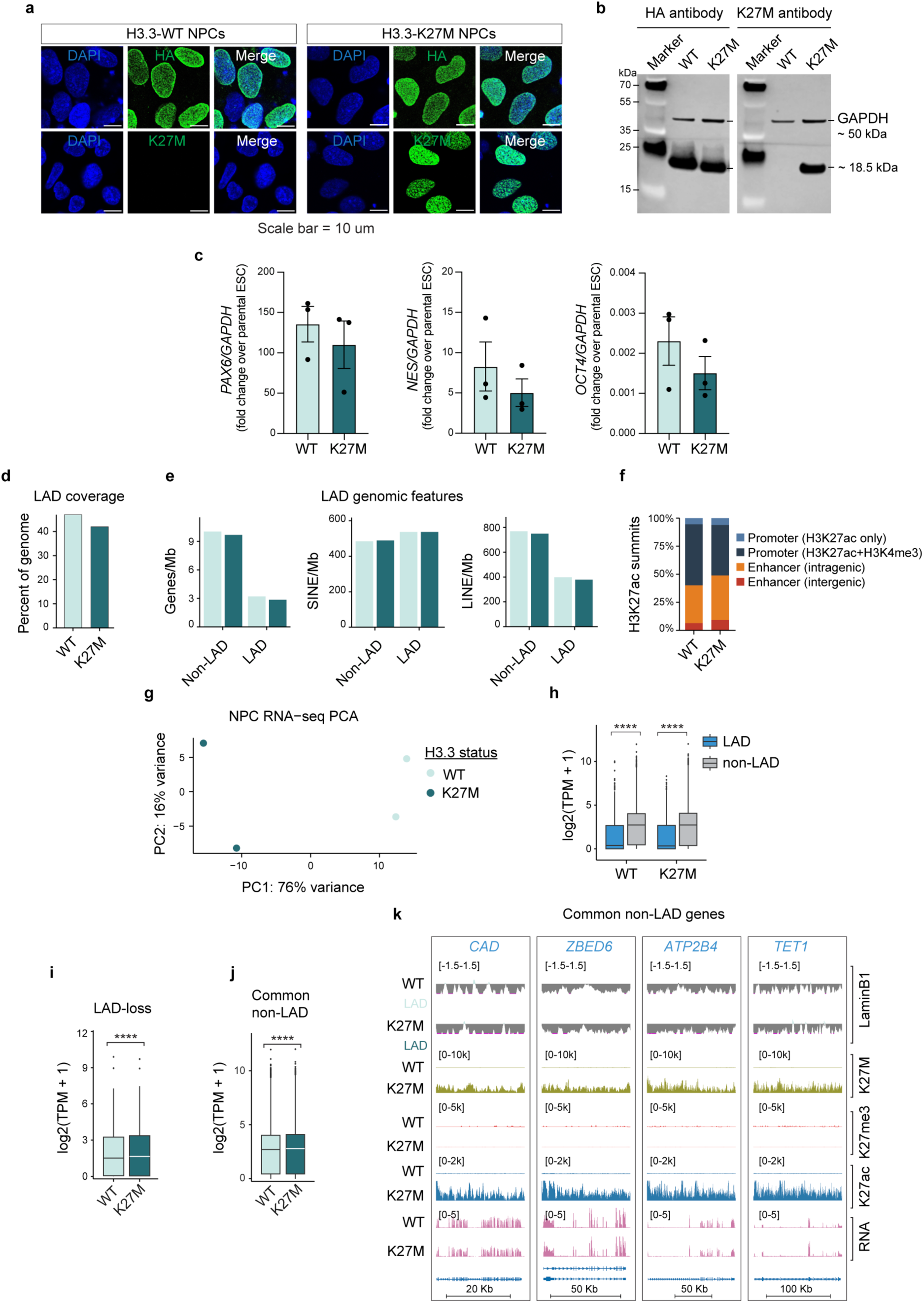
Validation of H3.3 expressing NPCs. A, Representative immunofluorescence images of ESCs after 3 days of doxycycline induction expressing H3.3WT-HA or H3.3K27M-HA. HA and H3K27M staining are shown in green and nuclear DNA is shown in blue. **B,** Western blot of H3.3WT and H3.3K27M NPCs against HA and H3K27M with GAPDH loading control. **C,** Expression of NPC marker genes *PAX6* and *NES* and ESC marker gene *OCT4* in H3.3WT and H3.3K27M NPCs relative to parental ESCs. **D,** Percentage of analyzed genome in LADs for H3.3WT and H3.3K27M NPCs. **E,** Gene, SINE, and LINE density in count per Mb in Non-LADs and LADs by genotype (light green = H3.3WT, dark green = H3.3K27M). **F,** Location of H3K27ac summits in H3.3 engineered NPCs. Promoters were defined as summits within 1 kb of annotated TSS ± presence of H3K4me3 peak. Enhancer summits were defined as all other peaks stratified by location within or outside annotated gene body. **G,** PCA of RNA-seq from engineered H3.3 NPCs with 2 replicates per genotype. **H,** Boxplots of gene expression shown as log2(TPM + 1) of LAD and non-LAD genes in H3.3 engineered NPCs. **I, J,** Boxplots of replicate averaged gene expression shown as log2(TPM + 1) of genes in LAD-loss regions (**I**) or Common non-LAD regions (**J**) for H3.3 engineered NPCs. Significance was assessed using unpaired Wilcoxon rank-sum tests. ****: P < 0.0001. **K,** Representative genome browser tracks of common non-LAD loci *CAD, ZBED6, ATP2B4,* and *TET1*. LaminB1 GO-CaRT signal is shown as log2(LMNB1/IgG). H3K27M, H3K27me3, and H3K27ac CUT&RUN signal are shown as spike-in normalized coverage signal. RNA is shown as bins per million.

